# Macrophage Iron Metabolism in Allografts and Tumors

**DOI:** 10.64898/2026.02.24.707843

**Authors:** Xiaohan Li, Xi Zhang, Ran Li, Tangchun Wu, Li Zhang, Zheng Gan, Yongjun Wang, Weicong Ye, Song Wang, Yanglin Hao, Kexiao Zheng, Zifeng Zou, Yinghuan Liu, Yilong Li, Zetong Tao, Jie Wu, Jiahong Xia

**Author notes:** These authors contributed equally to this work (Xiaohan Li, Xi Zhang, Ran Li).

## Abstract

Macrophages are present at high frequencies in transplanted organs and solid tumors. These infiltrating macrophages are highly tuned by tissue niche signals. However, the underlying mechanisms that orchestrate the activation of distinct macrophage populations remain unclear. Here, using heart transplantation and tumor implantation models, we found that macrophages in cardiac allografts exhibited higher intracellular iron (Fe^2+^) level and higher expression of the iron transporter SLC11A1 than those in the solid tumors. In mice, the myeloid-specific deletion of *Slc11a1* alleviated the proinflammatory state of macrophages and cardiac allografts rejection. Together, our findings identify SLC11A1 as a new player in immune response regulation, implicating SLC11A1 as a therapeutic target for both allograft and tumor rejection. This preprint reports initial findings; additional experiments are ongoing and will be included in a future full manuscript.

## INTRODUCTION

Macrophages are highly plastic and adapt their functions to environmental cues, with activation *in vivo* occurring in complex microenvironments that generate diverse functional statuses^1–5^. In tumors, macrophages predominantly adopt an immunosuppressive phenotype, suppressing tumor rejection through the upregulation of inhibitory checkpoints (PD-1, PD-L1, and Tim-3) and reducing the expression of MHCII in tumors, thereby suppressing antitumor immunity^6^. In contrast, in transplanted organs, they exhibit an immunostimulatory phenotype, promoting allograft rejection through increased expression of costimulatory molecules (CD80 and CD86) and MHCII and the secretion of proinflammatory cytokines (IL-1β, IL-6, and IL-12)^7,8^. The functional polarization of macrophages depends on the cytokine milieu, but accumulating evidence indicates that the metabolic context also plays a critical role in this process^9,10^. Solid tumors typically impose metabolic constraints characterized by nutrient (glucose, amino acids, lipids, and ions) deprivation, hypoxia, and acidosis, whereas transplanted organs provide a nutrient-rich and oxygen-sufficient environment^11,12^. Whether these distinct metabolic conditions drive the differential polarization of macrophages in tumors and transplanted organs remains unclear.

Iron is a ubiquitous ionic metabolite and plays a central role in both the differentiation and polarization of macrophages^13–15^. Tissue-infiltrating macrophages can sense and react to iron signaling of environmental cues to assist bystander parenchymal cells in their functional outputs and contribute to the immune response or suppression^16^. In spinal cord injury, iron accumulation increases TNF expression, resulting in persistent M1-like polarization that is detrimental to recovery^17^. Iron released from erythrophagocytosis in chronic venous leg ulcers drives a proinflammatory macrophage population, promoting persistent TNF-α-mediated inflammation, oxidative and nitrative stress, and fibroblast senescence, ultimately impairing wound healing^18^. However, in solid tumors, tumor cells upregulate lipocalin-2 (LCN2) and its receptor SCL22A17 in response to macrophage-derived cytokines to compete for limited iron levels, leading to iron depletion in macrophages^19^. This process leads to M2-like macrophage polarization and promotes tumor progression. Notably, the dietary commensal microbe *Lactiplantibacillus plantarum* IMB19 can drive macrophages to limit iron availability to tumor cells and adopt an iron-dependent proinflammatory phenotype, thereby suppressing tumor progression^20^. However, the role of iron metabolism in opposing macrophage functional polarization in transplanted organs and solid tumors, as well as the underlying mechanisms, are poorly understood.

In this study, we showed that SLC11A1 is a major contributor to the opposing macrophage functional polarization in transplanted organs and solid tumors. The myeloid-specific deletion of *Slc11a1* alleviates heart transplant rejection and the proinflammatory state of macrophages. Taken together, these results suggest that targeting SLC11A1 in macrophages is a therapeutic strategy for transplant rejection and cancer.

## RESULTS

### Distinct immune phenotypes and iron metabolic states in allograft and tumor infiltrating macrophages

The immune phenotype of immune cells is shaped by the tissue microenvironmental cues^21–23^. To characterize immune cell states in allografts and tumors, we established a murine B16-F10 melanoma model and performed single-cell RNA sequencing (scRNA-seq) of tumor-infiltrating CD45□ cells at day 20 after tumor implantation. This dataset was integrated with a publicly available scRNA-seq dataset of murine cardiac allografts harvested at day 7 post-transplantation. (https://figshare.com/s/11d8b3c99165c5fd8cd3) (Figure 1A). Differential expression analysis across all immune cell types between allografts and tumors revealed that macrophages exhibited the largest proportion of differentially expressed genes (Figure 1B). Based on this observation, we reclustered macrophages from the integrated dataset and identified two transcriptionally distinct clusters corresponding to allograft-infiltrating macrophages [Macs (Allograft)] and tumor-infiltrating macrophages [Macs (Tumor)] (Figure 1C). Macs (Allograft) exhibited significantly higher transplant rejection scores compared with Macs (Tumor) (Figure 1D). Further differential gene expression analysis between Macs (Allograft) and Macs (Tumor) demonstrated enrichment of pathways associated with macrophage functions and immune responses (Figure 1E). Metabolic programs were closely associated with the phenotypic and functional remodeling of macrophages^24–27^. Our data showed that Macs (Allograft) exhibited higher metabolism scores than Macs (Tumor) (Figure 1F). Given that metabolite transporters serve as key regulators of intracellular metabolic flux^28–30^, we profiled differentially expressed metabolite transporter genes between Macs (Allograft) and Macs (Tumor). We found that most differentially expressed transporters were associated with ion transport, with significant enrichment in iron ion transmembrane transport pathways (Figure 1G). Notably, Macs (Allograft) exhibited significantly higher iron ion transport scores than Macs (Tumor) (Figure 1H) and iron transport activity was strongly positively correlated with transplant rejection scores across both macrophage subsets (Figure 1I). To determine whether iron transport activity was associated with altered intracellular iron (Fe^2^□) levels, we performed flow cytometry analysis and found that macs (Allograft) exhibited higher intracellular iron (Fe^2^□) levels compared with macs (Tumor) (Figure 1J). Thus, these findings suggest that allograft- and tumor-infiltrating macrophages exhibit distinct immune phenotypes and iron metabolic states.

**Figure 1.**
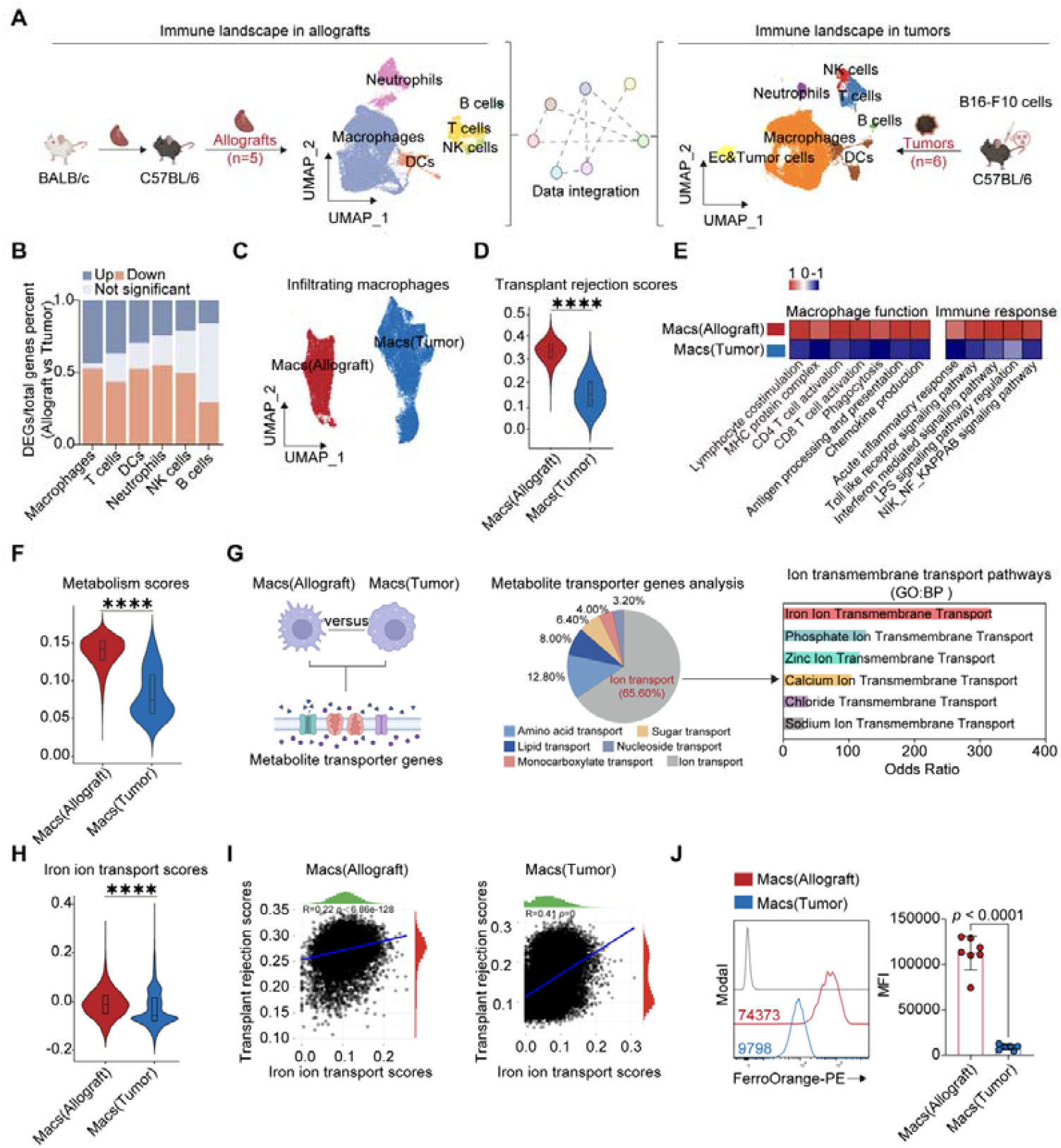
Distinct immune phenotypes and iron metabolic states in allograft and tumor infiltrating macrophages. (A) Schematic of the integrated strategy for scRNA-seq data from allografts and tumors. (B) The proportion of differentially expressed genes of each cell cluster. (C) UMAP plot showing the clustering of macrophages in allografts and tumors. (D) Transplant rejection scores (HALLMARK_ALLOGRAFT_REJECTION) of Macs (Allograft) and Macs (Tumor). (E) Heatmap showing the GSVA scores for the indicated pathways of Macs (Allograft) and Macs (Tumor). (F) Metabolism scores (REACTOME_METABOLISM) of Macs (Allograft) and Macs (Tumor). (G) Left panel, schematic of the comparative strategy for analyzing differentially expressed metabolite transporter genes between Macs (Allograft) and Macs (Tumor). Mid panel, functional categorization of differentially expressed metabolite transporter genes, including genes related to amino acid, sugar, lipid, nucleoside, monocarboxylate, and ion transport. Right panel, enriched GO: BP terms for ion transport genes ranked by odds ratio. (H) Iron ion transport scores (GO:0034755) of the Macs (Allograft) and Macs (Tumor). (I) Correlations between the iron ion transport scores and transplant rejection scores in Macs (Allograft) and Macs (Tumor). (J) Representative FACS histograms and bar graphs of the fluorescence intensity of intracellular iron (Fe^2+^) levels in Macs (Allograft) and Macs (Tumor) (n = 7 mice per group). Macs (Allograft) were collected on day 7 posttransplantation. Macs (Tumor) were collected on day 20 post tumor implantation. *P<0.05, **P<0.01, ***P<0.001, and****P<0.0001. Data are presented as mean ± SD. For statistical analyses, the unpaired t test (J) was applied.

### Different expression of the iron transporter SLC11A1 in allograft and tumor infiltrating macrophages

To explore the mechanisms underlying the distinct iron metabolic states in allograft- and tumor-infiltrating macrophages, we compared iron transporter gene expression between Macs (Allograft) and Macs (Tumor). We identified 23 differentially expressed iron transporter genes, among which SLC11A1 showed the most pronounced differential expression and was significantly upregulated (Allograft) (Figure 2A-C). Subsequently, flow cytometry and immunofluorescence staining assay also confirmed this result (Figure 2D–E). To extend these findings to clinical settings, we reanalyzed a published scRNA-seq dataset of human allograft-infiltrating immune cells^31,32^, and found that SLC11A1 expression in macrophages was upregulated during allograft rejection (Figure 2F–G). Tumor immune responses shared transcriptional features with allograft rejection programs^33,34^. Moreover, macrophages represent a key cellular target of immune checkpoint blockade (ICB), with enhanced tumor-associated immune responses following ICB treatment^6,35,36^. To assess the association between macrophage SLC11A1 expression and ICB therapy, we analyzed an available human tumor scRNA-seq dataset ^37^ and observed that SLC11A1 expression in macrophages was significantly upregulated following ICB treatment (Fig. 2H–I). In summary, we identified differential expression of the iron transporter SLC11A1 between allograft- and tumor-infiltrating macrophages.

**Figure 2.**
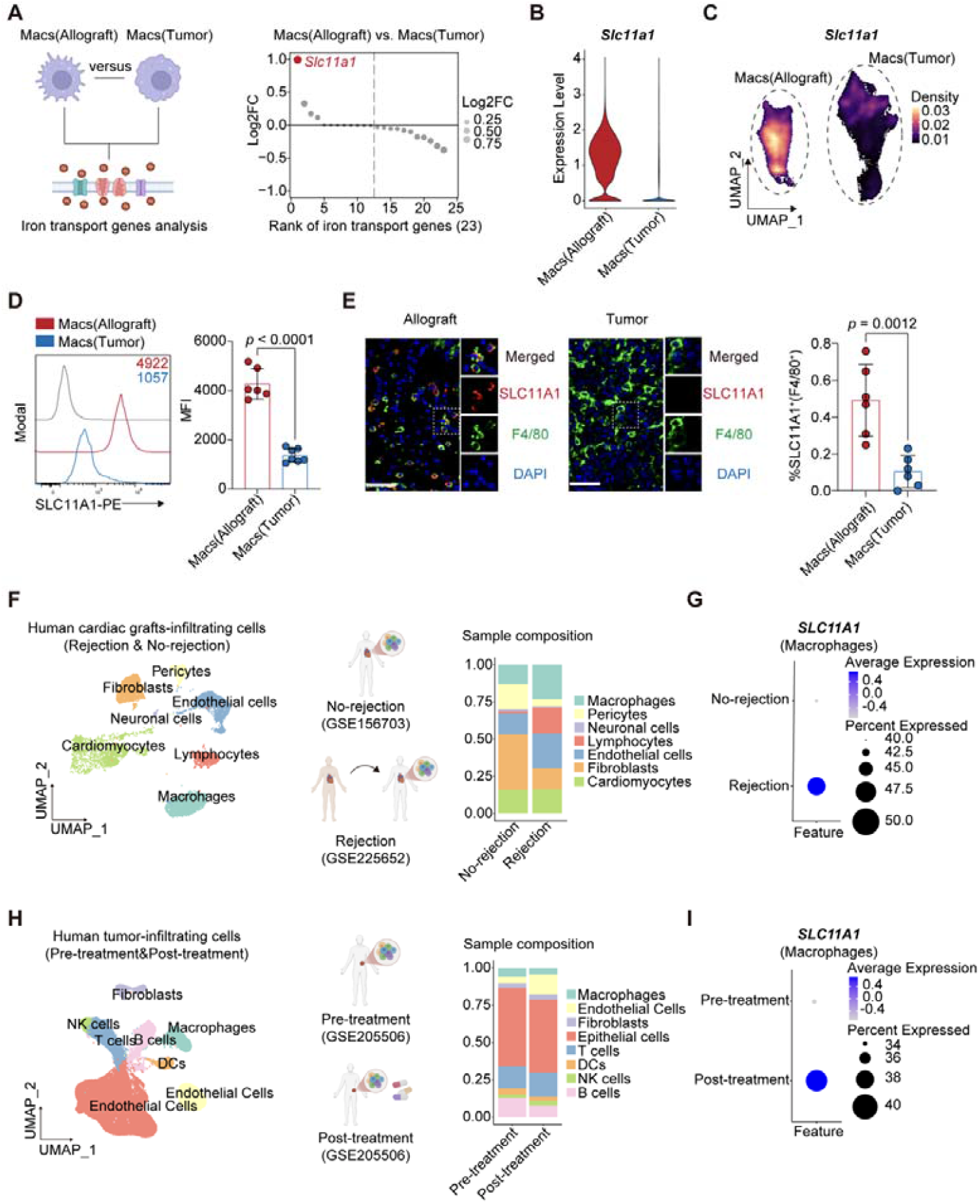
Different expression of the iron transporter SLC11A1 in allograft and tumor-infiltrating macrophages. (A) Left panel, schematic of the strategy used to identify differentially expressed iron transporter genes between Macs (Allograft) and Macs (Tumor). Right panel, differential expression of iron transporter genes between Macs (Allograft) and Macs (Tumor) ranked by log2FC values. (B) Violin plot showing *Slc11a1* expression in Macs (Allograft) and Macs (Tumor). (C) UMAP plot showing the expression intensity of *Slc11a1* in Macs (Allograft) and Macs (Tumor). (D) Representative FACS histograms and bar graphs of the fluorescence intensity values of SLC11A1 in Macs (Allograft) and Macs (Tumor) (n=7 mice per group). Macs (Allograft) were collected on day 7 posttransplantation. Macs (Tumor) were collected on day 20 post tumor implantation. (E) Representative images of immunofluorescence staining confirming SLC11A1 expression in allografts and tumors. Scale bars, 50 μm. Allografts were collected on day 7 posttransplantation. Tumors were collected on day 20 post tumor implantation. The bar graph shows the percentage of SLC11A1^+^(F4/80^+^) cells (n=6 mice per group). (F) UMAP plot and corresponding cell types proportions in human colorectal cancer scRNA-seq data (pre- and post-ICB treatment). (G) Dot plot showing *SLC11A1* expression in macrophage. (H) UMAP plot and corresponding cell types proportions of human allografts scRNA-seq data. (I) Dot plot showing *SLC11A1* expression in macrophage. Scale bars, 50 μm. Data are presented as mean ± SD. For statistical analyses, unpaired Student’s t test (D-E) was applied.

### Macrophage-specific *Slc11a1* deficiency alleviates allograft rejection

To investigate the role of SLC11A1 in macrophages, we generated myeloid-specific *Slc11a1*-conditional knockout mice (*Slc11a1*-cKO). Using a peritoneal macrophage activation model, allogeneic splenocytes were injected intraperitoneally (BALB/c to B6), and macrophages were harvested 16 h later for analysis (Figure 3A). Macrophage-specific deletion of *Slc11a1* significantly reduced the intracellular iron (Fe^2+^) levels (Figure 3B). Additionally, *Slc11a1* deficiency led to decreased expression of the costimulatory molecules (CD80 and CD86) and MHCII in macrophages (Figure 3B). To further investigate the role of SLC11A1 during allograft rejection, we established a full major histocompatibility complex (MHC) mismatch mouse heart transplantation model (Figure 3C). We found that *Slc11a1-*cKO mice prolonged allograft survival and attenuated inflammatory cell infiltration and myocyte injury (Figure 3D-E). Flow cytometric analysis showed that *Slc11a1-*cKO significantly decreased intracellular iron (Fe^2+^) levels in allograft-infiltrating macrophages (Figure 3F). This reduction was accompanied by the marked downregulation of the expression of CD80, CD86, MHCII, and the classical proinflammatory marker Ly6C (Figure 3F). As a major immune infiltrate in allografts, macrophages modulated effector T cell responses during allograft rejection^38,39^. T cells are key effector cells in allograft rejection, and their production of IFN-γ and TNF-α contributes to the rejection response^40–42^. Our result showed that *Slc11a1-*deficient mice exhibited reduced proportions and numbers of IFN-γ and TNF-α-producing CD4^+^ and CD8^+^ T cells compared with *WT* mice (Figure 3G-H). Taken together, these data indicate that macrophage-specific *Slc11a1* deficiency alleviates allograft rejection in mice and suggest that *Slc11a1* is a potential therapeutic target in allograft rejection.

**Figure 3.**
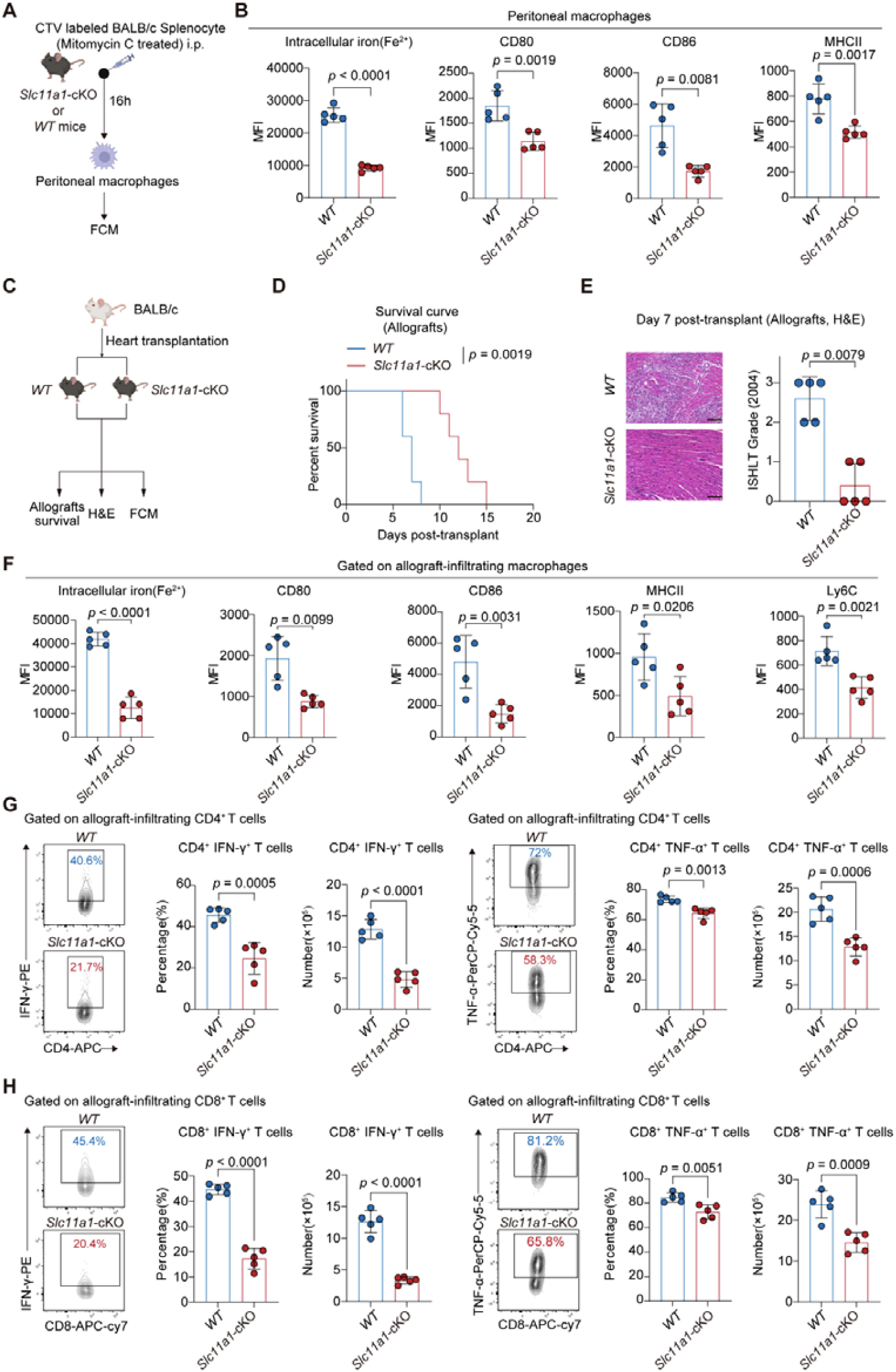
Macrophage-specific *Slc11a1* deficiency alleviates allograft rejection. (A) Schematic of the experimental design: mitomycin C-treated BALB/c splenocytes were labeled with or without CellTrace Violet (CTV) and intraperitoneally injected into *WT* (*Slc11a1*^fl/fl^) or *Slc11a1*cKO (*Slc11a1*^fl/fl^ *Lyz2*^cre+^) B6 mice. After 16 hours, the phagocytic capacity, intracellular iron (Fe^2+^) levels, and expression of costimulatory molecules, MHCII, and inflammatory markers in peritoneal macrophages were assessed by flow cytometry. The remaining peritoneal macrophages were cocultured with CTV-labeled TEa cells for 48 hours to evaluate T-cell responses. (B) Bar graphs of the fluorescence intensity values of intracellular iron (Fe^2+^) levels and CD80, CD86, MHCII expression in peritoneal macrophages (n=5 mice per group). (C) Schematic of the murine heterotopic heart transplantation model. (D) Cardiac allograft survival in *WT* and *Slc11a1*cKO recipient mice (n=5 mice per group). (E) Representative images of hematoxylin and eosin staining of allografts from *WT* and *Slc11a1*cKO mice collected on day 7 posttransplantation. Scale bars, 100 μm. Cellular rejection was scored according to the 2004 International Society for Heart and Lung Transplantation (ISHLT) guidelines (n=5 mice per group). (F) Bar graphs of the fluorescence intensity values of intracellular iron (Fe^2+^) levels, CD80, CD86, MHCII, and Ly6C in allograft-infiltrating macrophages on day 7 posttransplantation (n=5 mice per group). (G) Representative FACS plots and bar graphs of the proportions and numbers of CD4^+^IFN-γ^+^ and CD4^+^TNF-α^+^ cells in the spleen on day 7 posttransplantation (n=5 mice per group). (H) Representative FACS plots and bar graphs of the proportions and numbers of CD8^+^IFN-γ^+^ and CD8^+^TNF-α^+^ cells in the spleen on day 7 posttransplantation (n=5 mice per group). Data are presented as mean ± SD. For statistical analyses, the log-rank test (D), unpaired Student’s t test (B, F, G, H) and Mann□Whitney U test (E) was applied.

## DISCUSSION

In this study, we showed that the SLC11A1 drives opposing macrophage polarization in transplanted organs and tumors. The myeloid-specific deletion of *Slc11a1* alleviates heart transplant rejection and the proinflammatory state of macrophages. Therefore, targeting SLC11A1 in macrophages may be a novel therapeutic strategy for transplant rejection and cancer.

Tissue-infiltrating macrophages display substantial phenotypic and functional heterogeneity, driven by tissue niche signals, including metabolites^1^. Numerous studies have linked macrophage heterogeneity to the pathogenesis of multiple diseases. Maurizio et al. demonstrated that macrophages exhibit high plasticity in autoimmune and inflammatory diseases and contribute to tissue-specific pathology^43^. We identified two transcriptionally and functionally distinct macrophage subsets: allograft-infiltrating macrophages [Macs (Allograft)] and tumor-infiltrating macrophages [Macs (Tumor)]. These populations exhibit pronounced heterogeneity in their inflammatory response, cytokine production, and antigen presentation. Macs (Allograft) have the highest allograft rejection scores, whereas Macs (Tumor) have the lowest scores. This functional heterogeneity highlights the influence of the tissue microenvironment in determining macrophage immunophenotypic profiles.

Emerging evidence highlights that ion metabolism, particularly iron metabolism, is a critical regulator of immune cell function^14,15,44–46^. For instance, iron has been shown to control CD4^+^ T-cell effector activity by modulating proteins involved in glycolysis, RNA processing, and epigenetic regulation^47–49^. Moreover, Yuhang et al. reported that iron availability influences B-cell proliferation and antibody production through epigenetic mechanisms^50^. We revealed that the differentially expressed metabolite transporters between Macs (Allograft) and Macs (Tumor) are predominantly involved in ion transport and are significantly enriched in iron ion transport pathways. Macs (Allograft) presented elevated iron ion transport activity accompanied by increased intracellular iron (Fe^2+^) levels. Building on these findings, our study revealed that *Slc11a1*, a phagosomal iron transporter previously implicated in host defense against pathogens^51,52^, is a critical regulator of iron homeostasis in macrophages. SLC11A1 expression is selectively upregulated in Macs (Allograft) but is substantially repressed in Macs (Tumor). Moreover, macrophages extensively infiltrate allografts during transplant rejection and contribute to the progression of rejection through multiple mechanisms, such as antigen presentation and the production of inflammatory cytokines^7,8^. Using *Slc11a1*-conditional knockout mice, we observed that the deletion of *Slc11a1* significantly prolonged cardiac graft survival and ameliorated pathological scores. Macs (Allograft) exhibited decreased intracellular iron (Fe^2+^) levels and downregulated the expression of costimulatory molecules (CD80 and CD86), MHCII and the classical inflammatory marker Ly6C. In parallel, the proportion and number of IFN-γ^+^ and TNF-α^+^ CD4^+^ T cells were markedly reduced.

## ACKNOWLEDGMENTS

This work was supported by the National Natural Science Foundation of China [82422036, 82530061, 82271811, 82241217, 82572044], National Key Research and Development Program of China [2023YFC2706200], Natural science fund of Hubei Province [2024AFA047, 2025AFB477], the Fundamental Research Funds for the Central Universities [YCJJ20252116], the China Postdoctoral Science Foundation [2024T170310, 2023M741285]. The figures in this article were created with BioRender.com.

## EXPERIMENTAL MODEL AND SUBJECT DETAILS

### Mice

BALB/c (H-2d) and C57BL/6 (B6, H-2b) mice were purchased from BIONT Biological Technology (Hubei, China). *Slc11a1*^fl/fl^ (C57BL/6JGpt-*Slc11a1*^em1Cflox/Gpt^) mice were purchased from GemPharmatech (Jiangsu, China). *Lyz2-*Cre mice (C57BL/6 background) were purchased from the Jackson Laboratory (Maine, USA). Macrophage-specific *Slc11a1* gene knockout (*Slc11a1*^fl/fl^ Lyz2-Cre^+^) mice were generated by crossing *Slc11a1*^fl/fl^ mice with *Lyz2-*Cre mice. TEa (B6. Cg-Tg(Tcra, Tcrb)3Ayr/J) mice were purchased from Jackson Laboratory (Maine, USA). All the mice were aged 8 weeks, weighed 20–30 g and were housed in a specific pathogen-free laboratory at Huazhong University of Science and Technology (Wuhan, China). The mice used in our study were male, except when indicated otherwise. Ethical approval for the animal experiments was obtained from the Institutional Animal Care and Use Committee of Huazhong University of Science and Technology.

## METHOD DETAILS

### Murine heterotopic heart transplantation

Heterotopic heart transplantation in mice was performed in our laboratory as previously described^53^. Briefly, hearts were isolated from the donor mice, and the recipient mice were anesthetized. The donor heart was subsequently transplanted to the recipient, with the abdominal aortal and inferior vena caval incisions of the recipient anastomosed to the cardiac aorta and pulmonary artery of the donor. The function of the cardiac grafts was monitored daily by abdominal palpation. HTR was defined as the cessation of a palpable heartbeat and was confirmed by autopsy. In CTLA-4-treated heart transplantation models, after heterotopic heart transplantation, recipients were treated with cytotoxic T-lymphocyte-associated protein 4-immunoglobulin (CTLA4-Ig; Bristol-Myers-Squibb, 250 μg ip) on days 1 and 2.

### Tumor models

Syngeneic subcutaneous tumor models were established by resuspending 1×10^6^ B16-F10 tumor cells in 100□µL of PBS and injecting this mixture subcutaneously into the right flank of each mouse. Tumor growth was monitored every two days using digital calipers. Animals were euthanized for signs of distress or when the total tumor volume reached 1500 mm^3^.

### Cell staining and flow cytometry

Equal numbers of cells were stained for each sample, washed with ice-cold PBS and then blocked with an anti-CD16/32 antibody (BioLegend, 101302) on ice for 10 min before staining. The cell suspensions were first stained with a Zombie Aqua Fixable Viability Kit (BioLegend, 423102) for 15 min at room temperature. The surface proteins were then stained in FACS buffer (PBS containing 2% FBS and 0.1% sodium azide) for 30 min at 4 °C. For transcription factor staining, cells were fixed in Foxp3/Transcription Factor Staining Kit (Invitrogen, 00-5521-00) for 30 min at 4 °C and washed with 1× Permeabilization buffer (Invitrogen, 00-8333-56), followed by intracellular antibody staining for 30 min at 4 °C. Fe^2+^ levels were detected by staining the cells with FerroOrange (1□µM, MCE, HY-D1913) at 37□□ for 30□min. The CellTrace Violet (CTV) Kit (Thermo Fisher Scientific, C34557) was employed to assess T cell proliferation. To detect intracellular cytokines, cells were stimulated with 50 ng/ml phorbol 12-myristate 13-acetate (PMA, Abmole, M4647) and 500 ng/ml ionomycin (Sigma-Aldrich, I0634), and treated with GolgiStop (BD Biosciences, 554724) for 6 h following the manufacturer’s instructions. Cells were then fixed with BD Cytofix/Cytoperm (BD Biosciences, 554714) for 30 min at 4 °C, followed by washing with 1× permeabilization buffer. For indirect intracellular staining of SLC11A1 cells were fixed and permeabilized as described above and then stained with an SLC11A1 polyclonal antibody (Invitrogen, PA5-114373) at 37 °C for 30 min, followed by staining with a PE-conjugated donkey anti-rabbit IgG antibody (Biolegend, 406421) at 37 °C for 30 min.

### Histology and Immunofluorescence analysis

The harvested cardiac grafts were embedded in paraffin and processed into tissue sections. Subsequently, hematoxylin and eosin (H&E) staining was performed, and rejection response was scored in accordance with the 2004 International Society for Heart and Lung Transplantation (ISHLT) grading guidelines. Paraffin-embedded heart graft tissues were sectioned at a thickness of 5□μm and deparaffinized in xylene for 20 min. The sections were then rehydrated through a graded ethanol series (100%, 95%, 85%, 75%, and 50%) at room temperature. Antigen retrieval was performed by boiling the sections in retrieval buffer for 10 min, followed by three washes with PBS. The sections were subsequently permeabilized and blocked in PBS containing 0.3% Triton X-100 and 3% bovine serum albumin (BSA) for 1 hour at room temperature. Next, the sections were incubated overnight at 4□°C with primary antibodies. After three washes with PBS, the sections were incubated with isotype-matched secondary antibodies for 1 hour at room temperature. The nuclei were counterstained with 2□μg/mL DAPI for 10 min in the dark. Finally, the sections were mounted with antifade mounting medium, and fluorescence images were acquired using an OLYMPUS FV3000 confocal laser scanning microscope (Olympus Corporation, Japan).

### Isolation of infiltrating immune cells from cardiac grafts and tumors

Cardiac grafts were collected from recipient mice on day 7 posttransplantation for the isolation of graft-infiltrating immune cells. Briefly, graft tissues were minced into small fragments and enzymatically digested with 1□mg/mL collagenase B (Roche, 11088815001) in Hank’s balanced salt solution containing calcium and magnesium (Solarbio, H1025) at 37□°C for 20 min. The digestion mixture was collected every 5 min, gently resuspended, and transferred to prechilled tubes on ice. Following enzymatic digestion, the resulting cell suspension was passed through 70-μm cell strainers to remove debris. Mononuclear cells were then enriched by density gradient centrifugation using 40% Percoll (Solarbio, P8370) at 600 × g for 25 min at 25□°C without applying a brake. After centrifugation, the mononuclear cell layer was collected, washed, resuspended in PBS, and then transferred to flow cytometry tubes for surface marker staining and analysis. Tumor tissues were collected and processed using a Mouse Tumor Dissociation Kit (Miltenyi Biotec, 130-096-730) according to the manufacturer’s instructions. Briefly, excised tumors were minced into small fragments, enzymatically digested with the enzyme mixture at 37□°C for 30 min, and mechanically dissociated. The cell suspension was filtered through a 70-μm cell strainer in RPMI-1640 medium. The cells were then centrifuged at 300 × g for 3 min and further purified by Percoll gradient centrifugation.

### Isolation of peritoneal macrophages (PMs)

Splenocytes were isolated from BALB/c mice and labeled ex vivo with 2.0□µM CellTrace Violet. Following CTV labeling, the cells were treated with mitomycin C (Stemcell, 73274) and injected into the peritoneal cavity of C57BL/6 or *Slc11a1*^fl/fl^ *Lyz2*-Cre^+^ mice. Sixteen hours after the injection, the host mice were euthanized, and the peritoneal cavity was flushed three times consecutively with 9□mL of cold DPBS. After 1 min, the peritoneal lavage fluid was collected by centrifugation at 1700 rpm for 5 min. The phagocytic activity (measured by uptake of CTV^+^ cells) and activation status of macrophages were assessed by flow cytometry.

### Single-cell RNA sequencing (scRNA-seq) and data analysis

The single-cell suspensions used for the scRNA-seq analysis were prepared as described above. The scRNA-seq expression matrices were processed using Seurat (version 4.0.1), which provides tools for quality control, filtering, normalization, and dimensionality reduction. Low-quality cells were removed based on the following criteria: cells with fewer than 500 or more than 5000 detected genes or a mitochondrial gene content exceeding 5%. Genes expressed in at least five cells were retained for downstream analysis. The ‘ScaleData’ function was applied to center and scale the integrated data. PCA was performed using the ‘RunPCA’ function, and the first 20 principal components were used for dimensionality reduction and clustering via the ‘FindClusters’ function. UMAP was used to visualize the resulting clusters. Marker genes for each cluster were identified using the ‘FindAllMarkers’ function. Cell types were annotated based on a combination of automated classification tools and a comparison with previously published high-throughput datasets. Violin plots of gene expression were generated using the ‘VlnPlot’ function. Gene set scores were computed using Seurat’s ‘Add Module Score’ function, AUCell or GSVA package, which calculates the average expression of each predefined gene set at the single-cell level. These scores were visualized in violin plots.

## QUANTIFICATION AND STATISTICAL ANALYSIS

GraphPad Prism 10.9.0 was used for statistical analyses. The results are presented as the means mean ± SD, unless indicated otherwise. Unpaired Student’s t tests were used for comparisons between two groups when the data fit a normal distribution and had similar variances. Unpaired t test with Welch’s correction was used for normally distributed data when the variances were different, and the Mann□Whitney U test was used for nonnormally distributed data. Differences between multiple groups were assessed by one-way ANOVA or two-way ANOVA for normally distributed data followed by Dunnett’s, Sidak or Tukey multiple comparisons test. For Kaplan-Meier survival analysis, the log-rank test was used.

## DATA AVAILABILITY AND CODE AVAILABILITY

The data that support the findings of this study are available from the corresponding author upon reasonable request.

## REFERENCES

1. Guan, F., Wang, R., Yi, Z., Luo, P., Liu, W., Xie, Y., Liu, Z., Xia, Z., Zhang, H., and Cheng, Q. (2025). Tissue macrophages: origin, heterogenity, biological functions, diseases and therapeutic targets. Sig Transduct Target Ther 10, 93. 10.1038/s41392-025-02124-y.

2. Okabe, Y., and Medzhitov, R. (2016). Tissue biology perspective on macrophages. Nat Immunol 17, 9–17. 10.1038/ni.3320.

3. Epelman, S., Lavine, K.J., and Randolph, G.J. (2014). Origin and Functions of Tissue Macrophages. Immunity 41, 21–35. 10.1016/j.immuni.2014.06.013.

4. Locati, M., Curtale, G., and Mantovani, A. (2020). Diversity, Mechanisms, and Significance of Macrophage Plasticity. Annu. Rev. Pathol. Mech. Dis. 15, 123–147. 10.1146/annurev-pathmechdis-012418-012718.

5. Sheu, K.M., and Hoffmann, A. (2022). Functional Hallmarks of Healthy Macrophage Responses: Their Regulatory Basis and Disease Relevance. Annu. Rev. Immunol. 40, 295–321. 10.1146/annurev-immunol-101320-031555.

6. Christofides, A., Strauss, L., Yeo, A., Cao, C., Charest, A., and Boussiotis, V.A. (2022). The complex role of tumor-infiltrating macrophages. Nat Immunol 23, 1148–1156. 10.1038/s41590-022-01267-2.

7. Owen, M.C., and Kopecky, B.J. (2024). Targeting Macrophages in Organ Transplantation: A Step Toward Personalized Medicine. Transplantation 108, 2045–2056. 10.1097/TP.0000000000004978.

8. Panzer, S.E. (2022). Macrophages in Transplantation: A Matter of Plasticity, Polarization, and Diversity. Transplantation 106, 257–267. 10.1097/TP.0000000000003804.

9. Wculek, S.K., Dunphy, G., Heras-Murillo, I., Mastrangelo, A., and Sancho, D. (2022). Metabolism of tissue macrophages in homeostasis and pathology. Cell Mol Immunol 19, 384–408. 10.1038/s41423-021-00791-9.

10. Wculek, S.K., Forisch, S., Miguel, V., and Sancho, D. (2024). Metabolic homeostasis of tissue macrophages across the lifespan. Trends in Endocrinology & Metabolism 35, 793–808. 10.1016/j.tem.2024.04.017.

11. Lv, Y., Li, Z., Liu, S., Zhou, Z., Song, J., Ba, Y., Weng, S., Zuo, A., Xu, H., Luo, P., et al. (2025). Metabolic checkpoints in immune cell reprogramming: rewiring immunometabolism for cancer therapy. Mol Cancer 24, 210. 10.1186/s12943-025-02407-6.

12. Tao, Z., Luo, Z., Zou, Z., Ye, W., Hao, Y., Li, X., Zheng, K., Wu, J., Xia, J., Zhao, Y., et al. (2025). Novel insights and an updated review of metabolic syndrome in immune-mediated organ transplant rejection. Front. Immunol. 16, 1580369. 10.3389/fimmu.2025.1580369.

13. Frost, J.N., and Drakesmith, H. (2025). Iron and the immune system. Nat Rev Immunol. 10.1038/s41577-025-01193-y.

14. Ludwig, N., Cucinelli, S., Hametner, S., Muckenthaler, M.U., and Schirmer, L. (2024). Iron scavenging and myeloid cell polarization. Trends in Immunology 45, 625–638. 10.1016/j.it.2024.06.006.

15. Mu, Q. The role of iron homeostasis in remodeling immune function and regulating inflammatory disease.

16. Soares, M.P., and Hamza, I. (2016). Macrophages and Iron Metabolism. Immunity 44, 492–504. 10.1016/j.immuni.2016.02.016.

17. Kroner, A., Greenhalgh, A.D., Zarruk, J.G., Passos dos Santos, R., Gaestel, M., and David, S. (2014). TNF and Increased Intracellular Iron Alter Macrophage Polarization to a Detrimental M1 Phenotype in the Injured Spinal Cord. Neuron 83, 1098–1116. 10.1016/j.neuron.2014.07.027.

18. Sindrilaru, A., Peters, T., Wieschalka, S., Baican, C., Baican, A., Peter, H., Hainzl, A., Schatz, S., Qi, Y., Schlecht, A., et al. (2011). An unrestrained proinflammatory M1 macrophage population induced by iron impairs wound healing in humans and mice. J. Clin. Invest. 121, 985–997. 10.1172/JCI44490.

19. Zhang, Y.-Y., Han, Y., Li, W.-N., Xu, R.-H., and Ju, H.-Q. (2024). Tumor iron homeostasis and immune regulation. Trends in Pharmacological Sciences 45, 145–156. 10.1016/j.tips.2023.12.003.

20. Sharma, G., Sharma, A., Kim, I., Cha, D.G., Kim, S., Park, E.S., Noh, J.G., Lee, J., Ku, J.H., Choi, Y.H., et al. (2024). A dietary commensal microbe enhances antitumor immunity by activating tumor macrophages to sequester iron. Nat Immunol 25, 790–801. 10.1038/s41590-024-01816-x.

21. Ren, X., Zhang, L., Zhang, Y., Li, Z., Siemers, N., and Zhang, Z. (2021). Insights Gained from Single-Cell Analysis of Immune Cells in the Tumor Microenvironment. Annu. Rev. Immunol. 39, 583–609. 10.1146/annurev-immunol-110519-071134.

22. Pearce, E.L., and Pearce, E.J. (2013). Metabolic Pathways in Immune Cell Activation and Quiescence. Immunity 38, 633–643. 10.1016/j.immuni.2013.04.005.

23. Park, J., Hsueh, P.-C., Li, Z., and Ho, P.-C. (2023). Microenvironment-driven metabolic adaptations guiding CD8+ T cell anti-tumor immunity. Immunity 56, 32–42. 10.1016/j.immuni.2022.12.008.

24. Yan, J., and Horng, T. (2020). Lipid Metabolism in Regulation of Macrophage Functions. Trends in Cell Biology 30, 979–989. 10.1016/j.tcb.2020.09.006.

25. Koelwyn, G.J., Corr, E.M., Erbay, E., and Moore, K.J. (2018). Regulation of macrophage immunometabolism in atherosclerosis. Nat Immunol 19, 526–537. 10.1038/s41590-018-0113-3.

26. Wculek, S.K., Dunphy, G., Heras-Murillo, I., Mastrangelo, A., and Sancho, D. (2022). Metabolism of tissue macrophages in homeostasis and pathology. Cell Mol Immunol 19, 384–408. 10.1038/s41423-021-00791-9.

27. Jin, R., Neufeld, L., and McGaha, T.L. (2025). Linking macrophage metabolism to function in the tumor microenvironment. Nat Cancer 6, 239–252. 10.1038/s43018-025-00909-2.

28. Lin, L., Yee, S.W., Kim, R.B., and Giacomini, K.M. (2015). SLC transporters as therapeutic targets: emerging opportunities. Nat Rev Drug Discov 14, 543–560. 10.1038/nrd4626.

29. Reina-Campos, M., Scharping, N.E., and Goldrath, A.W. (2021). CD8+ T cell metabolism in infection and cancer. Nat Rev Immunol 21, 718–738. 10.1038/s41577-021-00537-8.

30. Packer, M. (2023). SGLT2 inhibitors: role in protective reprogramming of cardiac nutrient transport and metabolism. Nat Rev Cardiol 20, 443–462. 10.1038/s41569-022-00824-4.

31. Li, X., Turaga, D., Li, R.G., Tsai, C.-R., Quinn, J.N., Zhao, Y., Wilson, R., Carlson, K., Wang, J., Spinner, J.A., et al. (2024). The Macrophage Landscape Across the Lifespan of a Human Cardiac Allograft. Circulation 149, 1650–1666. 10.1161/CIRCULATIONAHA.123.065294.

32. Sim, C.B., Phipson, B., Ziemann, M., Rafehi, H., Mills, R.J., Watt, K.I., Abu-Bonsrah, K.D., Kalathur, R.K.R., Voges, H.K., Dinh, D.T., et al. (2021). Sex-Specific Control of Human Heart Maturation by the Progesterone Receptor. Circulation 143, 1614–1628. 10.1161/CIRCULATIONAHA.120.051921.

33. Dunn, G.P., Bruce, A.T., Ikeda, H., Old, L.J., and Schreiber, R.D. (2002). Cancer immunoediting: from immunosurveillance to tumor escape. Nat Immunol 3, 991–998. 10.1038/ni1102-991.

34. Schreiber, R.D., Old, L.J., and Smyth, M.J. (2011). Cancer Immunoediting: Integrating Immunity’s Roles in Cancer Suppression and Promotion. Science 331, 1565–1570. 10.1126/science.1203486.

35. Zhang, H., Liu, L., Liu, J., Dang, P., Hu, S., Yuan, W., Sun, Z., Liu, Y., and Wang, C. (2023). Roles of tumor-associated macrophages in anti-PD-1/PD-L1 immunotherapy for solid cancers. Mol Cancer 22, 58. 10.1186/s12943-023-01725-x.

36. Cassetta, L., and Pollard, J.W. (2023). A timeline of tumour-associated macrophage biology. Nat Rev Cancer 23, 238–257. 10.1038/s41568-022-00547-1.

37. Li, J., Wu, C., Hu, H., Qin, G., Wu, X., Bai, F., Zhang, J., Cai, Y., Huang, Y., Wang, C., et al. (2023). Remodeling of the immune and stromal cell compartment by PD-1 blockade in mismatch repair-deficient colorectal cancer. Cancer Cell 41, 1152–1169.e7. 10.1016/j.ccell.2023.04.011.

38. Wyburn, K.R., Jose, M.D., Wu, H., Atkins, R.C., and Chadban, S.J. (2005). The Role of Macrophages in Allograft Rejection. Transplantation 80, 1641–1647. 10.1097/01.tp.0000173903.26886.20.

39. Jose, M.D., Ikezumi, Y., Van Rooijen, N., Atkins, R.C., and Chadban, S.J. (2003). Macrophages act as effectors of tissue damage in acute renal allograft rejection. Transplantation 76, 1015–1022. 10.1097/01.TP.0000083507.67995.13.

40. Koh, K.P., Wang, Y., Yi, T., Shiao, S.L., Lorber, M.I., Sessa, W.C., Tellides, G., and Pober, J.S. (2004). T cell–mediated vascular dysfunction of human allografts results from IFN-γ dysregulation of NO synthase. J. Clin. Invest. 114, 846–856. 10.1172/JCI21767.

41. Räisänen-Sokolowski, A., Mottram, P.L., Glysing-Jensen, T., Satoskar, A., and Russell, M.E. (1997). Heart transplants in interferon-gamma, interleukin 4, and interleukin 10 knockout mice. Recipient environment alters graft rejection. J. Clin. Invest. 100, 2449–2456. 10.1172/JCI119787.

42. Azzawi, M. (1999). Tumour necrosis factor alpha and the cardiovascular system: its role in cardiac allograft rejection and heart disease. Cardiovascular Research 43, 850–859. 10.1016/S0008-6363(99)00138-8.

43. Cutolo, M., Soldano, S., Smith, V., Gotelli, E., and Hysa, E. (2025). Dynamic macrophage phenotypes in autoimmune and inflammatory rheumatic diseases. Nat Rev Rheumatol 21, 546–565. 10.1038/s41584-025-01279-w.

44. Rodriguez, R., Müller, S., Colombeau, L., Solier, S., Sindikubwabo, F., and Cañeque, T. (2025). Metal Ion Signaling in Biomedicine. Chem. Rev. 125, 660–744. 10.1021/acs.chemrev.4c00577.

45. Marques, O., Weiss, G., and Muckenthaler, M.U. (2022). The role of iron in chronic inflammatory diseases: from mechanisms to treatment options in anemia of inflammation. Blood 140, 2011–2023. 10.1182/blood.2021013472.

46. Marques, O., Weiss, G., and Muckenthaler, M.U. (2022). The role of iron in chronic inflammatory diseases: from mechanisms to treatment options in anemia of inflammation. Blood 140, 2011–2023. 10.1182/blood.2021013472.

47. Wang, Z., Yin, W., Zhu, L., Li, J., Yao, Y., Chen, F., Sun, M., Zhang, J., Shen, N., Song, Y., et al. (2018). Iron Drives T Helper Cell Pathogenicity by Promoting RNA-Binding Protein PCBP1-Mediated Proinflammatory Cytokine Production. Immunity 49, 80–92.e7. 10.1016/j.immuni.2018.05.008.

48. Lai, Y., Zhao, S., Chen, B., Huang, Y., Guo, C., Li, M., Ye, B., Wang, S., Zhang, H., and Yang, N. (2022). Iron controls T helper cell pathogenicity by promoting glucose metabolism in autoimmune myopathy. Clinical & Translational Med 12, e999. 10.1002/ctm2.999.

49. Gao, X., Song, Y., Wu, J., Lu, S., Min, X., Liu, L., Hu, L., Zheng, M., Du, P., Yu, Y., et al. (2022). Iron-dependent epigenetic modulation promotes pathogenic T cell differentiation in lupus. Journal of Clinical Investigation 132, e152345. 10.1172/JCI152345.

50. Jiang, Y., Li, C., Wu, Q., An, P., Huang, L., Wang, J., Chen, C., Chen, X., Zhang, F., Ma, L., et al. (2019). Iron-dependent histone 3 lysine 9 demethylation controls B cell proliferation and humoral immune responses. Nat Commun 10, 2935. 10.1038/s41467-019-11002-5.

51. Cunrath, O., and Bumann, D. (2019). Host resistance factor SLC11A1 restricts Salmonella growth through magnesium deprivation. Science 366, 995–999. 10.1126/science.aax7898.

52. Lokken-Toyli, K.L., Diaz-Ochoa, V.E., Camacho, L., Stull-Lane, A.R., Van Hecke, A.E.R., Mooney, J.P., Muñoz, A.D., Walker, G.T., Hampel, D., Jiang, X., et al. (2024). Vitamin A deficiency impairs neutrophil-mediated control of Salmonella via SLC11A1 in mice. Nat Microbiol 9, 727–736. 10.1038/s41564-024-01613-0.

53. Zhang, X., Xu, H., Yu, J., Cui, J., Chen, Z., Li, Y., Niu, Y., Wang, S., Ran, S., Zou, Y., et al. (2023). Immune Regulation of the Liver Through the PCSK9/CD36 Pathway During Heart Transplant Rejection. Circulation 148, 336–353. 10.1161/CIRCULATIONAHA.123.062788.

